# Restoring hippocampal sharp-wave ripples under glutamate transporter dysfunction

**DOI:** 10.64898/2026.05.28.728386

**Authors:** Yunhan Gao, Zeyu Zhou, Jian-young Wu

## Abstract

Disruption of glutamate homeostasis is believed to contribute to the early progression of Alzheimer’s disease (AD) and associated neurodegeneration. Soluble amyloid-β oligomers impair excitatory amino acid transporters (EAATs), reducing glutamate clearance, while also enhancing glutamate release from neurons and astrocytes. Together, these effects produce persistent glutamatergic dysregulation that disrupts synaptic and network function. Here, we asked whether the effects of EAAT attenuation can be mitigated through ion-channel modulation. TBOA, a selective EAAT inhibitor, was used to model early-stage glutamatergic dysregulation. TBOA reduced the local field potential amplitude of hippocampal sharp-wave ripples (SWRs) in mouse hippocampal slices, suggesting that glutamate accumulation disrupts network synchrony. Calcium imaging further showed that TBOA diminished SWR-associated population calcium transients while promoting spontaneous calcium transients in individual neurons, indicating a shift from coordinated population activity toward disorganized cellular activity. KCNQ-channel openers ML213 and ICA-27243 partially restored the TBOA-induced decline in SWR amplitude. In contrast, similar restorative effects were not observed following modulation of other ion channels, including blockade of AMPA and NMDA receptors or HCN/Ih channels, or activation of large-conductance Ca^2+^-activated K^+^ (BK) channels and G-protein–activated inwardly rectifying potassium (GIRK) channels. Together, these findings suggest that KCNQ-channel openers may occupy a unique position in mitigating glutamate-related hyperexcitability during early AD-associated network dysfunction.

## Introduction

Alzheimer’s disease (AD) is a progressive neurodegenerative disorder that undergoes decades of asymptomatic progression before the emergence of significant memory loss and cognitive decline [1]. Increasing evidence indicates that amyloid-beta (Aβ)–related alterations in neurotransmitter systems play a critical role in disease progression [2–7]. Among these systems, glutamatergic signaling—which underlies learning, memory, and synaptic plasticity—is particularly vulnerable during the preclinical stages of AD [8–10]. Dysregulation of glutamate disrupts neuronal communication and contributes to excitotoxicity and long-term neurodegeneration [11–16].

Mild elevation of extracellular glutamate is increasingly recognized as an important contributor to AD pathology [8, 9, 13, 17–19]. Soluble Aβ oligomers impair neuronal and astrocytic excitatory amino acid transporters (EAATs), thereby reducing glutamate clearance from the synaptic cleft [9], while simultaneously enhancing glutamate release from neurons and astrocytes. These combined effects produce persistent glutamatergic dysregulation that disrupts synaptic function and promotes network instability and neurodegeneration [20]. However, whether neuronal network dysfunction produced by mild but chronic elevations of extracellular glutamate can be reversed through compensatory mechanisms remains unknown.

To address this question, we modeled glutamate transporter dysfunction using DL-threo-β-benzyloxyaspartate (TBOA), a selective inhibitor of EAATs [21–22]. Previous studies have shown that ongoing neuronal activity is required for TBOA to induce hyperactivity, because continuous glutamate release allows extracellular glutamate to accumulate under conditions of EAAT attenuation [8]. In contrast, TBOA fails to induce hyperactivity in *ex vivo* cortical slices where spontaneous neuronal activity is minimal [23]. We therefore used hippocampal slices that naturally generate spontaneous network activity—sharp-wave ripples (SWRs)—providing an *ex vivo* preparation in which ongoing neuronal activity is preserved. This preparation allowed us to examine how TBOA-induced glutamate accumulation affects ongoing neuronal network dynamics and whether these effects can be mitigated through counteractive mechanisms.

SWRs are spontaneous neuronal population events that occur in the hippocampus during sleep and quiet wakefulness [24–27] and are associated with brain-wide activity involved in episodic memory consolidation [28–31]. These events can also be reliably observed in *ex vivo* hippocampal slices when cholinergic tone is low [25, 32]. In our slices, SWRs occur at a frequency of approximately 0.5–1 events per second, consistent with previous reports [23–25]. These robust SWRs therefore provide a sensitive probe for examining how mild glutamate elevation disrupts hippocampal network activity and how such disruption may be mitigated.

Here, we show that TBOA profoundly suppresses the amplitude of SWRs in hippocampal slices. Importantly, mild hyperpolarization produced by activation of voltage-gated potassium channels underlying the M-current restores SWR activity. These findings suggest that glutamate transporter dysfunction selectively disrupts organized hippocampal population events such as SWRs—network patterns associated with memory processing—while favoring increased spontaneous neuronal firing previously reported in other studies [17, 24]. In contrast, modest enhancement of potassium conductance stabilizes hippocampal network dynamics under conditions of glutamate transporter dysfunction. Our results further indicate that enhancing M-current–mediated membrane stabilization may represent a potential strategy for counteracting network dysfunction associated with early Aβ pathology.

## Methods

### Acute *ex vivo* brain slices

Hippocampal slices were prepared from P21–P75 male and female mice of the following genotypes: wild-type (C57BL/6), transgenic GCaMP6 [36] (C57BL/6J-Tg (Thy1-GCaMP6f), also known as GP5.5 Dkim/J, JAX #024276). We keep a homozygous colony in our institute (The Georgetown colony [35]), or transgenic ChR2 (B6.Cg-Tg(Thy1-COP4/EYFP)18Gfng/J, JAX #007612, WuLab homozygous colony).

All procedures were conducted in accordance with protocols approved by the Institutional Animal Care and Use Committee at Georgetown University Medical Center. Mice were deeply anesthetized with isoflurane and rapidly decapitated. Brains were quickly removed and chilled in ice-cold (0 °C) sucrose-based artificial cerebrospinal fluid (sACSF) containing (in mM): 252 sucrose, 3 KCl, 2 CaCl₂, 6 MgSO₄, 1.25 NaH₂PO₄, 26 NaHCO₃, and 10 dextrose, continuously bubbled with 95% O₂/5% CO₂.

Obliquely horizontal hippocampal slices (480 μm thick) were sectioned at a 12.7° fronto-occipital angle [37-38, Figure 1B] using a vibratome (Leica VT1000S) or compresstome (VF-300, Precisionary Instruments). Slices were incubated in artificial cerebral spinal fluid (ACSF) containing (in mM): 132 NaCl, 3 KCl, 2 CaCl₂, 2 MgSO₄, 1.25 NaH₂PO₄, 26 NaHCO₃, and 10 dextrose, bubbled with 95% O₂/5% CO₂, at 30–32 °C for 30 minutes, then gradually cooled to 26–27 °C for recovery over 3–7 hours.

**Fig 1.**
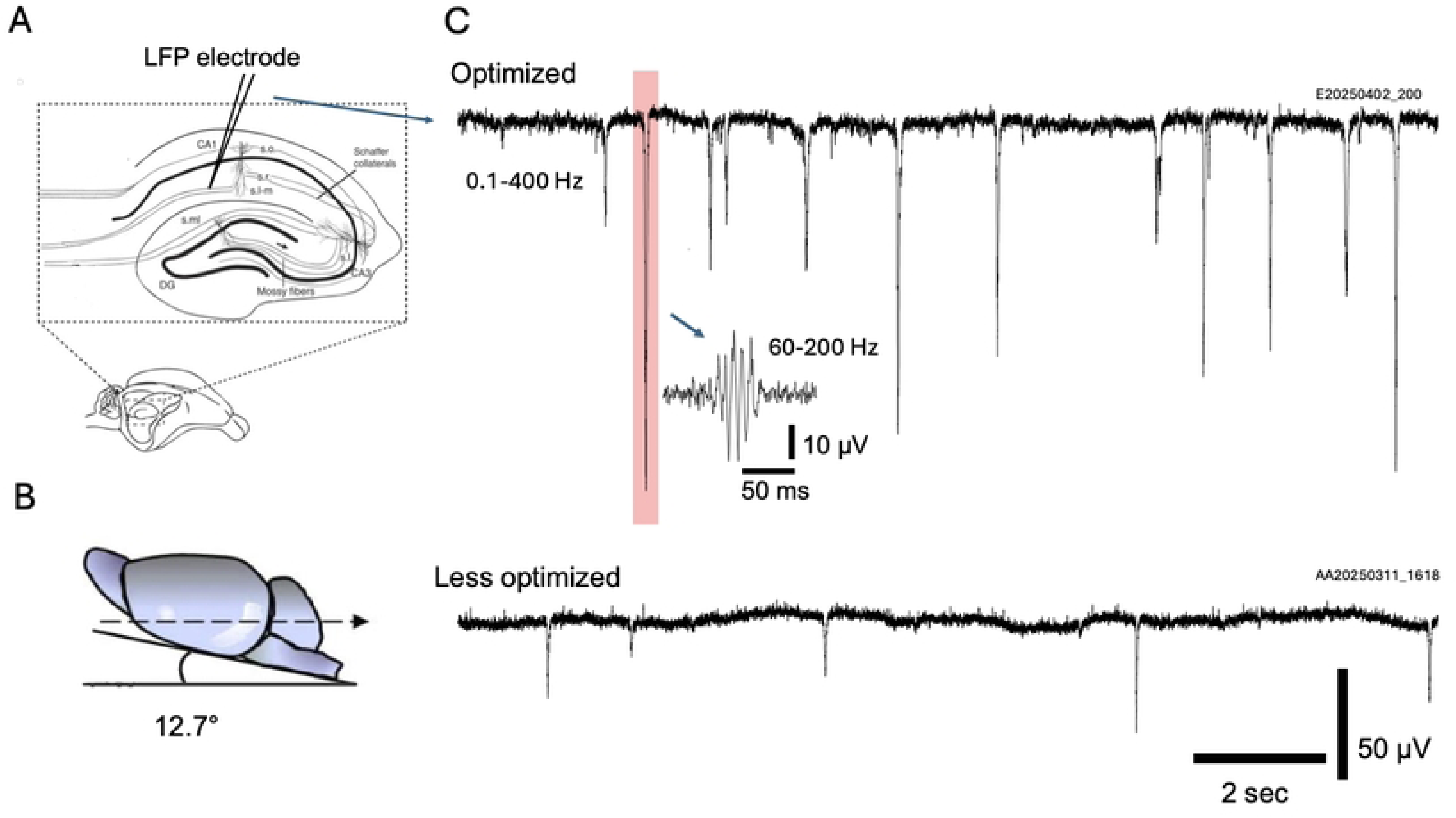
Reliable recording of SWRs. **A.** Schematic of the LFP recording configuration. SWRs were recorded from the stratum radiatum of the ventral hippocampus. **B.** An oblique slicing angle of 12.7° was used to preserve Schaffer collateral connectivity. **C.** Representative LFP traces bandpass filtered at 0.1– 400 Hz. Top trace: an optimized slice(in cutting angle, reduced dragging force during slicing and adequate recovery time) showing frequent, high-amplitude SWRs. *Inset:* a single SWR (shaded) filtered at 60–200 Hz to reveal ripple oscillations. Bottom trace: a less optimal slice showing infrequent, lower-amplitude SWRs.

### Local field potential (LFP) recording

LFP recordings were done in a custom submerged chamber, and slices were placed on a mesh that allowed ACSF perfusion on both sides at a high flow rate (10–30 ml /min) [39–40]. We used a custom ultra-low noise recording electrode and amplifier system, which can reliably distinguish low amplitude SWR events (∼5 μV). Low resistance glass microelectrodes (∼50 kΩ tip resistance) were pulled with a Sutter P87 puller with 6–7 controlled pulls. The electrode was filled by capillary action when the tip was in contact with the ACSF. Using ACSF as an intrapipette solution prevents potential alteration of the ionic condition in the tissue surrounding the electrode tip. The recording electrode was placed in CA1 stratum radiatum (Fig 1A). In each slice, several locations were tried to find an optimum recording location with high signal-to-noise. Usually, the highest SWR peak was >100 µV, and the ripple amplitude was 2–3 times higher than the noise level. To reduce 60Hz noise, a reference electrode was placed near the recording electrode, outside of the tissue. The LFP signal was amplified 1000x with a custom-made probe amplifier, filtered at 0.01–1000Hz and digitized at 3000 Hz by a 12-bit USB Analog-to-digital converter (National instruments). From each brain slice, we recorded 3–18 hours of spontaneous SWRs from the same recording site.

### Calcium imaging

GCaMP6f fluorescence signals were recorded using a cooled CCD camera (NeuroCCD 80×80). Signals were expressed as ΔF/F [41], defined as (F_x_ − F_0_)/F_0_, where F_x_ is the fluorescence signal from each region of interest (ROI) and F_0_ is the baseline fluorescence of the same ROI. Images were acquired at 125 frames/s using Turbo-SM64 software (SciMeasure/RedShirt Imaging). Imaging was performed on an upright microscope (Olympus BX51WI) with 470 nm LED excitation (Thorlabs) through a GFP filter cube (Chroma; excitation 425–475 nm, dichroic 480 nm, emission >485 nm). A 10X objective (0.30 NA, Amscope) was used to image large areas of the hippocampal CA3 and CA1 subfields. Illumination intensity at the sample was <10 mW/mm², under which photobleaching was negligible, allowing 2000–6000 s of imaging per field of view with adequate signal-to-noise ratio. Imaging was divided into consecutive trials (100 s per trial), with typically 30–80 trials collected per slice.

### Processing of Ca²⁺ imaging data

Each recording trial (100 s) consisted of 12,500 images. Raw images were processed using Turbo-SM64 or custom Python software to generate two data representations: signal traces and image frames. Traces quantify fluorescence changes (ΔF) at individual pixels or ROIs (defined as the linear average of ΔF across selected pixels), expressed as ΔF/F to reflect calcium transient amplitude [41]. Image frames depict the spatial extent of events, with pixel intensities normalized to show relative fluorescence across the field of view.

To generate peak images for each SWR event, custom software was used to automatically identify the peak time. Two averaged images were then constructed: a peak image (mean of 20 frames; 10 before and 10 after the peak) and a background image (mean of 30 frames acquired 100 ms before the peak). The SWR image was obtained by subtracting the background image from the peak image and normalizing pixel values between the minimum and maximum intensities, yielding a 128-level grayscale image representing the spatial extent of each SWR event.

### Experimental design and statistical analyses

Most experiments in this study have been presented in a case-control experimental design, in which the effects of drugs are compared to the control conditions. In analyzing the impact of factors on LFP activity in a large sample of *ex vivo* slices, we have taken a factorial design. All data were tested for normality and lognormality via Shapiro-Wilk tests. If all groups were normally distributed, they were analyzed with parametric tests (unpaired t-test, n-way ANOVA, n-way RM ANOVA) and were displayed on bar plots with error bars representing the mean ± SEM. Data were tested for non-uniformity via a Rayleigh test to determine if they could be sampled from a von Mises (circular normal) distribution. If the Rayleigh test reached significance for all groups, means were compared with the Watson-Williams test, otherwise a circular median test was performed.

The particular statistical tests used are listed in the Results. Any values cited in the text are mean ± SEM. All post hoc multiple comparisons used the Šidák correction. Within each plot, all individual data points were presented. No data were excluded based on their values, but only for experimental reasons (e.g., no SWRs present, excessive slice movement, or electrode drifting). The n was indicated in the text and figure legends, and differs between experiments. Raw p-values were displayed in plots.

For calcium imaging data, significance was assessed using thresholding based on baseline noise (mean ± 2–3 SD) to identify ROI regions. When repeated measurements were obtained from the same slice, repeated-measures designs were used.

## Results

Local field potential (LFP) was recorded from CA1 stratum radiatum (str. rad.) in *ex vivo* slices (Fig 1A). Sharp wave peak amplitudes of 10–300 μV were commonly observed, with an occurrence rate of 0.5–3 events per second (Fig 1C). In each slice, the SWR amplitude varied across different locations within the str. rad., probably related to the preservation of intact Schafer Collateral fibers between CA3 and CA1. Before each experiment, we tested several str. rad. locations to find a recording location with high SWR peak amplitude. In most of our slices, the highest sharp wave peaks were >100 µV (Fig 1C), and the ripple amplitude was 2–3 times the noise level (Fig 1C, insert). Higher SWR amplitude suggests a larger fraction of neurons surrounding the electrode tip was viable and activated by each SWR generated from the CA3. A higher occurrence rate also suggests better connections between CA3 and CA1. Slicing at the 12.7° oblique angle (Fig 1B) [37–38] seemed to be crucial for viable SWRs. Low SWR amplitude and occurrence rate (Fig 1C, bottom trace) are often associated with imperfect slicing, where the vibratome blade dragged the tissue, causing changes in slicing angle and/or thickness. These slices were excluded from our results.

Efforts were made to obtain stable SWR amplitude and occurrence rate over a duration of 5–12 hours, which was essential for examining gradual changes caused by mild extracellular glutamate accumulation, which usually takes hours to occur.

Consistent with our previous reports [33–35]. Under control conditions, SWRs typically maintained stable occurrence rates and amplitudes for 2–3 hours, followed by a gradual decline during prolonged recordings.

### TBOA reduced SWR amplitude

Low concentration of DL-TBOA (6–16 μM) or TFB-TBOA (5–50 nM) induced a substantial reduction in SWR amplitude (Fig 2A). The TBOA concentrations were about ¼ to ½ of those that induce epileptiform activity [8], and did not induce epileptiform-like spikes or other hyperexcitable activity in most of our slices. The amplitude of SWR was gradually reduced when perfused at these concentrations (Fig 2B, E, red lines), suggesting a correlation with the accumulation of glutamate.

**Fig 2.**
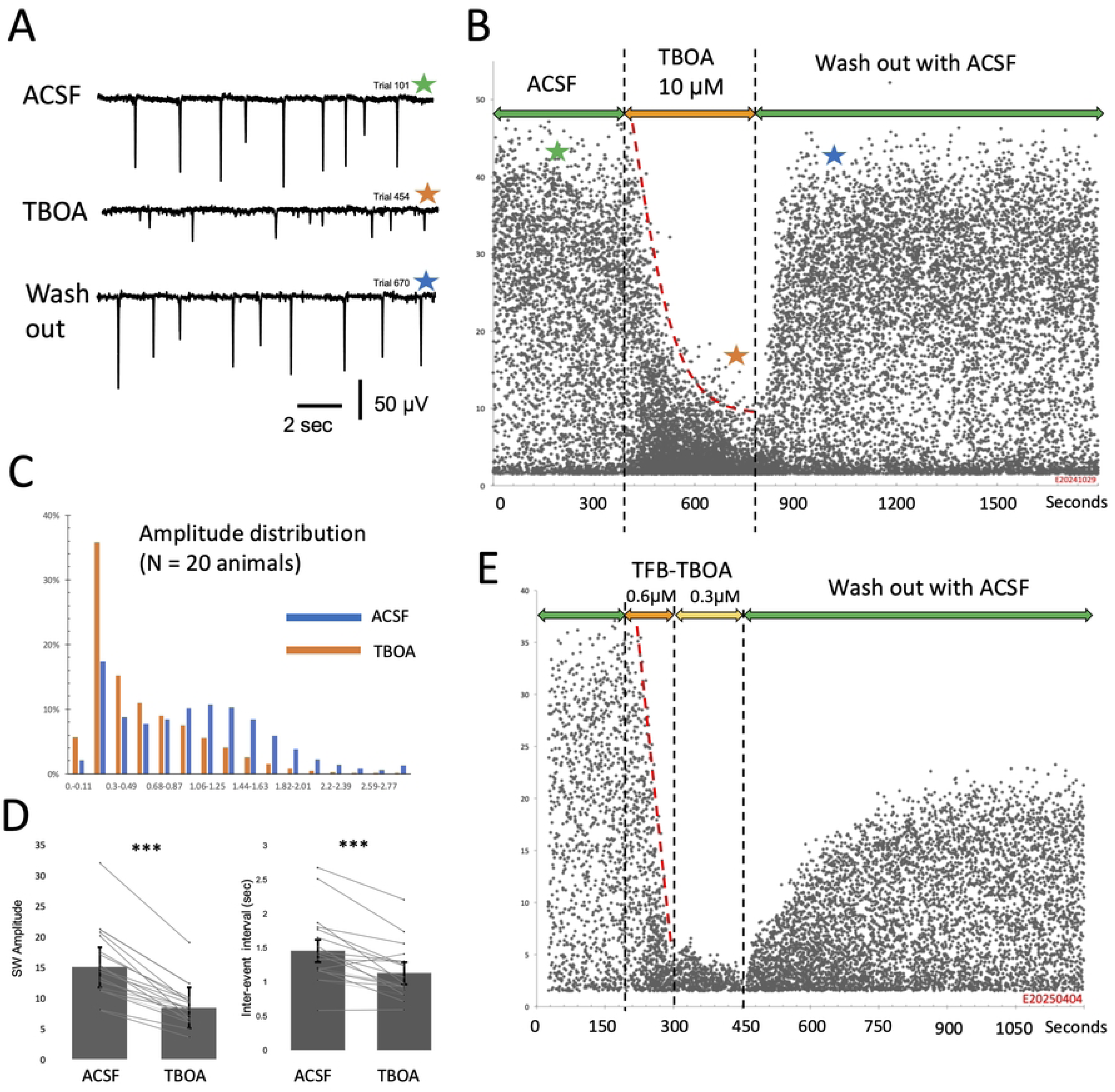
TBOA suppresses SWRs. **A.** Representative LFP traces of SWRs recorded under ACSF, TBOA, and washout conditions. Colored stars indicate the time points corresponding to traces shown in B. **B.** Time course of SWR amplitude from a representative experiment. Each dot represents the amplitude and time of an individual SWR event. TBOA (10 µM; shaded region between dashed lines) markedly suppresses SWR amplitude, which partially recovers after washout. The red dashed line indicates a smoothed trend of SWR amplitude over time. Colored stars indicate the time point corresponding to the traces shown in A. **C.** Distribution of SWR amplitudes pooled from 20 animals, showing a leftward shift toward smaller amplitudes in TBOA. **D.** Summary data from 20 animals showing mean SWR amplitude and inter-event interval (IEI). Each line represents an individual animal. TBOA significantly reduces SWR amplitude and increases IEI (***, p < 0.001; mean ± SEM). **E.** TFB-TBOA induced a stronger suppressive effect compared to TBOA, at lower concentrations (0.3–0.6 µM), with partial recovery after washout. Dashed lines indicate drug application periods; red dashed line show the smoothed amplitude trend.

Amplitude reduction caused a large shift in SWR amplitude distribution, in significantly more SWRs with low amplitude and much fewer high amplitude events (Fig 2C). TBOA-induced SWR amplitude reduction was seen in all slices tested (Fig 2D, left panel).

### TBOA increased SWRs occurrence rate

TBOA caused a small but statistically significant elevation in SWR occurrence rate, as indicated by a reduction in the inter-event intervals (Fig 2D, right panel) from 1.45 s (control average) to 1.12 s (TBOA average, paired two-tailed test; n=21 recordings; p=3.45×10⁻⁵), suggesting TBOA also affect the initiation of SWRs in the CA3 network.

### TBOA effects are reversible

SWR amplitude was quickly recovered after washing out of TBOA. This was obviously seen in raw recording traces (Fig 2A) and in accumulated data (Fig 2B, E). Higher TBOA concentration caused a more rapid decline of SWR amplitude and incomplete recovery (Fig 2E), suggesting that higher concentration also produces irreversible changes in the network. Occasional spontaneous hyperexcitable events were also seen, consistent with reports of other groups [8,42].

### TBOA disrupts organized SWR events and promotes disorganized activity

We next used calcium imaging to examine how TBOA disrupts organized SWR activity by inducing disorganized calcium fluctuations. In our Thy1 promoted GCaMP6f animals, a large fraction of CA1 pyramidal neurons is labeled, so calcium imaging can also see subthreshold calcium transients in the population in addition to that of spiking of individual neurons [43]. While calcium transients from individual neurons were poorly correlated with SWRs [44–45], “population” calcium transients have a 1:1 correlation to the SWRs in LFP recordings [43] (Fig 3B). Under control conditions, each SWR event evoked a population calcium signal diffusely distributed in the stratum radiatum of the CA1 region (Fig 3). The SWR-associated calcium signal was relatively small, with a ΔF/F of 0.1–2% (Fig 3B, trace). Spatially, SWR-associated calcium transients appeared in a broad area in the CA1 (Fig 3B, images).

**Fig 3.**
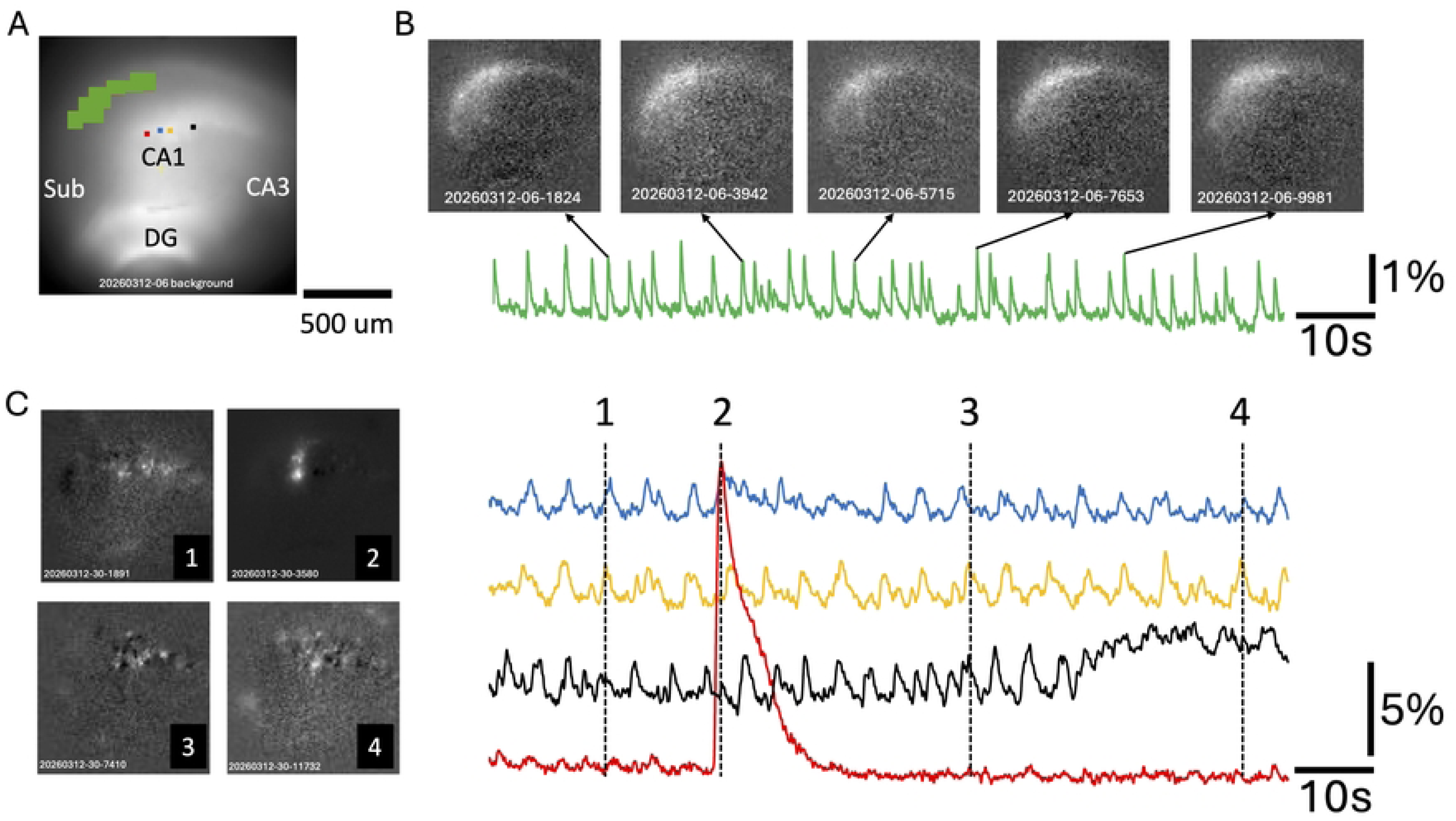
TBOA disrupts organized calcium activity during SWRs. **A.** Resting GCaMP6f fluorescence image of a hippocampal slice. The green ROI indicates the region used to extract population SWR-associated calcium signals (shown in B). Red, blue, orange, and black boxes are single pixel ROIs mark locations used to extract calcium transients from individual neurons under TBOA (traces shown in C). **B.** SWR-associated calcium activity under control conditions. The green trace represents the spatially integrated ΔF/F signal from the CA1 region (green ROI in A). Images are snapshots of ΔF/F at the peak of individual SWR events (time points indicated by arrows). ΔF/F was calculated as the change in fluorescence relative to baseline preceding each event. **C.** Neuronal activity under TBOA. Traces show ΔF/F signals from four individual pixels (ROIs in A), revealing large, localized calcium transients. Corresponding images (left) are snapshots (1-4) showing spatially restricted neuronal activity. The time of the snapshots are marked on the right traces (1-4). Note that a sustained Ca^2+^ elevation in the black trace between time 3-4.

Application of TBOA diminished the SWR-associated population calcium transients while promoting spontaneous calcium transients in individual neurons (Fig 3C). In the calcium imaging, “population” signals appear as distributed small calcium signals over a large area (Fig 3B images), while neuronal activity appears as localized “bright spots” (Fig 3C images). Three patterns of TBOA-associated calcium transients were seen: i) Repetitive transients peaks of similar amplitude (Fig 3C, blue, orange, and black traces), suggesting spontaneous firing of individual neurons. These individual peaks were not temporally aligned, suggesting asynchronous firing. ii) High amplitude transient signals (Fig 3C red trace), suggesting a high-frequency burst of a single neuron. iii) Sustained elevation of calcium signal (Fig 3C, right portion of the black trace), suggesting sustained firing. These imaging results indicate that TBOA exerts a differential effect on hippocampal network activity: it suppresses organized population events such as SWRs while promoting disorganized neuronal activity.

### Voltage-gated potassium channel openers reversed TBOA effects

TBOA effects of suppression of SWR and promoting unorganized firing were likely caused by a mild depolarization of the neuronal population due to extracellular glutamate. We hypothesize that such mild depolarization could be counteracted by a mild hyperpolarization produced through increased potassium conductance. To test this hypothesis, we applied potassium channel subfamily Q (KCNQ) openers to induce a small hyperpolarizing shift in the neuronal population.

We perfused slices with the KCNQ channel openers ML213 (KCNQ2/4)(1.5 µM) or ICA-27243 (KCNQ2/3)(0.62/1.5µM) after SWRs had been suppressed by TBOA and observed a pronounced reversal of the TBOA-associated SWR amplitude reduction (Fig 4A). SWR amplitude was significantly increased to near normal average. We also observed that the suppressive effect of TBOA accumulated over time; during continuous exposure to TBOA, SWR amplitude progressively declined even with the K openers.

**Fig 4.**
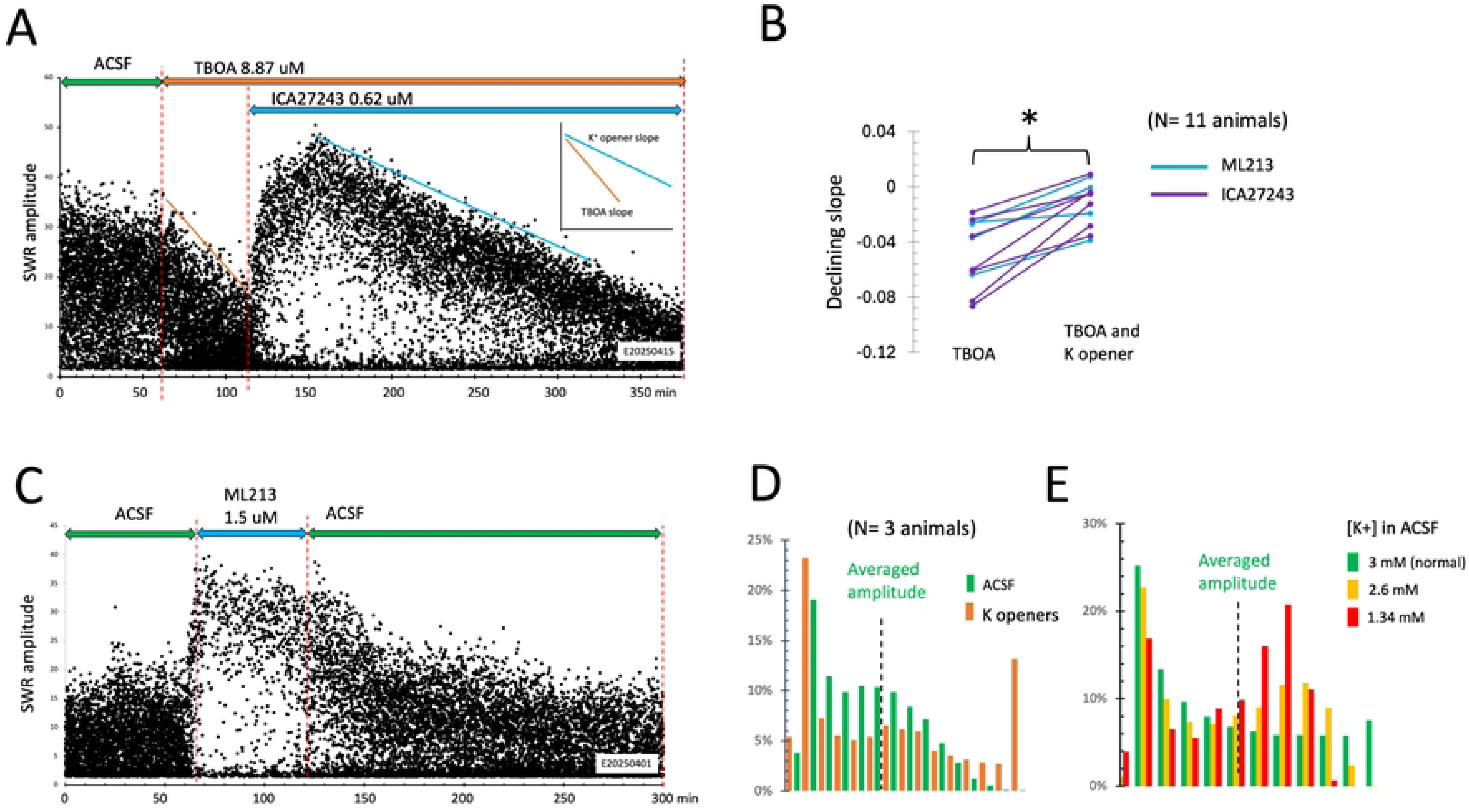
KNCQ openers reduced the declining rate of TBOA. **A.** SWR amplitude plots from a representative experiment under control, TBOA and KNCQ opener ICA27243. Double arrows mark the time of ACSF, TBOA or ICA27243 perfusion. Blue and orange lines mark the declining slope of TBOA only and TBOA+ IC27243. **B.** SWR amplitude declining slopes from 11 animals. **C.** KNCQ opener shifted the SWR amplitude distribution. **D - E.** SWR amplitude distribution in ACSF, KNCQ opener and low potassium ACSF.

However, while KCNQ openers could not reverse the TBOA effects, the progression of the SWR amplitude declining was significantly slowed, as seen in the altered declining slope (Fig 4B). Notably, K+ openers also changed SWR amplitude (Fig 4C) and amplitude distributions (Fig 4C). These changes are in the opposite direction of TBOA, which induced smaller SWRs and a shift in the amplitude distribution to the lower end (Fig 2C), forming a “compression” in the amplitude-time plot (Fig 2B,E). In contrast, K+ openers induced higher SWR amplitude (Fig 4C) and shift the amplitude distribution to the higher end (Fig 4D). These results suggest that mild hyperpolarization of the neuronal population can enhance the synchronization of the CA1 neurons during SWR events.

KCNQ channel openers are expected to hyperpolarize neurons. We therefore asked whether a similar membrane-potential shift could mitigate TBOA-induced SWR suppression. To directly examine the potassium-dependent effect, we lowered the [K^+^] in the perfusing ACSF in the absence of TBOA or KCNQ openers. Lowering the potassium produced similar changes in SWR amplitude as well as a shift in SWR amplitude distribution (Fig 4E), and the effect was dose dependent, with an increased proportion of large-amplitude events and a reduced proportion of small ones.

Together, these results support the interpretation that potassium-dependent hyperpolarization counteracts the depolarizing effect of elevated extracellular glutamate and partially restores the neuronal population participating in each SWR event.

### Other potassium channel openers

We next tested whether Large-conductance Ca^2+^-activated K^+^ channels (BK) and G-protein–activated inwardly rectifying potassium channels (GIRK) channel openers can also rescue the TBOA suppression of SWR events. BK channels are large conductance (200–300 pS) potassium channels that are activated by both membrane depolarization and elevated intracellular Ca²⁺, providing a rapid repolarization to neurons after an action potential [46]. GIRK channels, which are widely involved in Gi/o-coupled receptors, such as GABA_B, opioid, dopamine, and serotonin 5-HT1A receptors, producing membrane hyperpolarization that slows neuronal firing and reduces excitability [47–48]. Both hyperpolarization mechanisms might counteract TBOA-induced SWR suppression.

BK channel openers BMS204352 (13.9 µM) or NS11021 (3.91 µM) and GIRK channel activator ML297(13.9 µM) produced no detectable change in SWR amplitude in control conditions and did not have a detectable effect on rescuing TBOA-induced suppression (data not shown). These results suggest that KCNQ channel openers are in a unique situation of rescuing TBOA induced SWR suppression.

### Glutamate channel antagonists cannot rescue TBOA effects

Glutamate receptor antagonists, which are widely investigated as strategies to reduce pathological hyperexcitability in neurological disorders, including Alzheimer’s disease and traumatic brain injury. Among them, NMDA receptor antagonists such as memantine were clinically used to limit excessive NMDA receptor activation, thereby reducing glutamate-mediated hyperexcitability and excitotoxic stress [49–51, 60]. We used NMDA receptor antagonist AP5 to test whether NMDA receptor antagonists can rescue TBOA induced SWR suppression. Applying AP5 at 100 μM can fully block NMDA receptor-mediated activity. [54–55] but did not show a significant change in SWR amplitude (Fig 5C, D), consistent with a previous report from another group [56]. AP5 did not slow down the amplitude reduction by TBOA (Fig 5D), demonstrating that blocking NMDA channels cannot reduce the TBOA induced SWR amplitude reduction.

**Fig 5.**
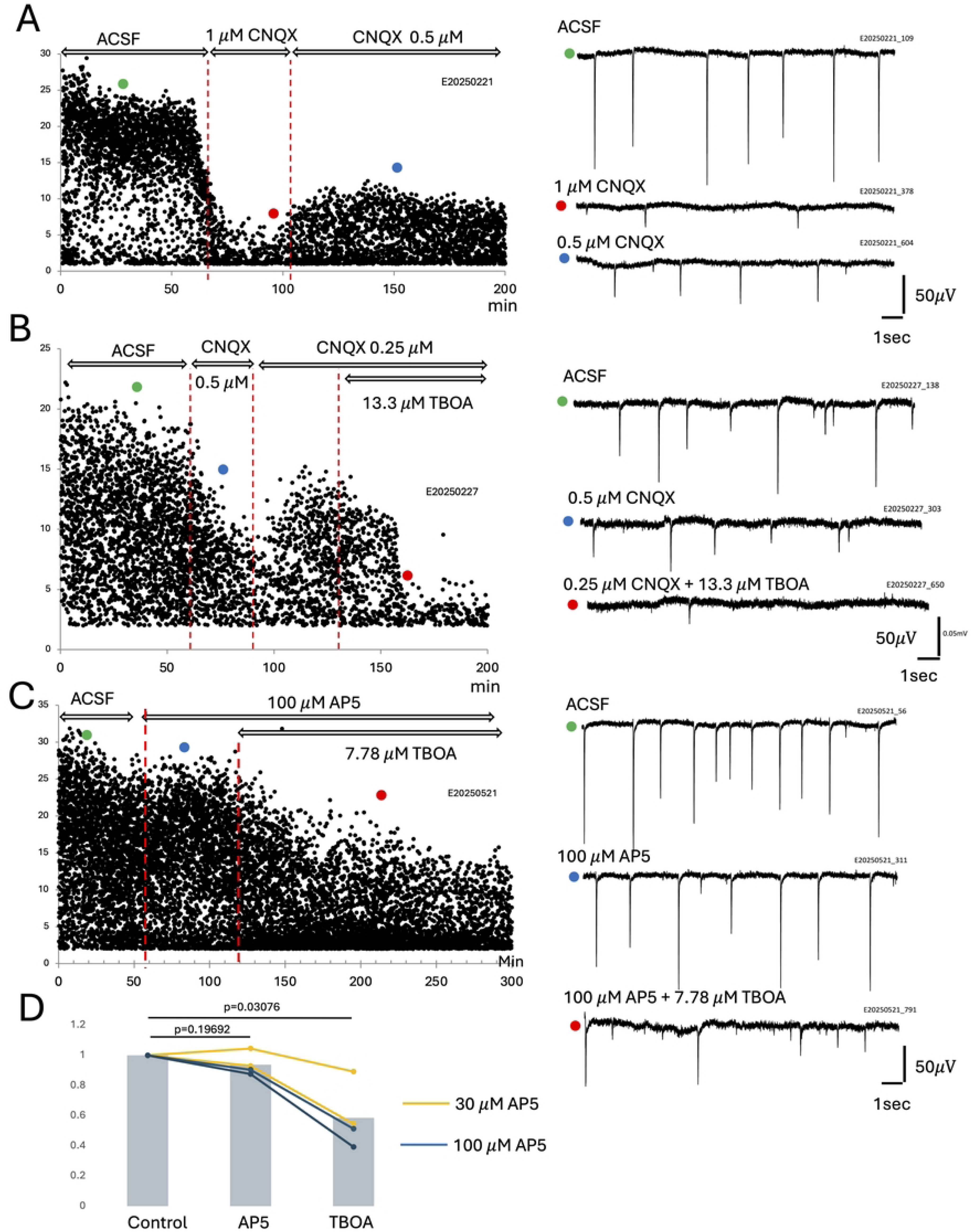
Glutamate receptor antagonists do not rescue TBOA-induced suppression of sharp waves. **A.** Low concentrations of CNQX suppressed SWR amplitude. Left, time course of SWR amplitude during perfusion with 1 μM and 0.5 μM CNQX. Right, representative recording traces under ACSF, 1 μM CNQX, and 0.5 μM CNQX. Colored dots on the left plot indicate the time points corresponding to the traces on the right. **B.** TBOA-induced suppression of SWRs was exacerbated in the presence of CNQX. Left, SWR amplitude plot during perfusion with 0.25–0.5 μM CNQX followed by 13.3 μM TBOA. Right, representative traces recorded under ACSF, 0.5 μM CNQX, and 0.25 μM CNQX + 13.3 μM TBOA. Note the sharp reduction in SWR amplitude when CNQX and TBOA were applied together. **C.** AP5 did not rescue TBOA-induced suppression of SWR amplitude. Left, SWR amplitude plot during perfusion with 100 μM AP5 followed by 7.78 μM TBOA. Right, representative traces under ACSF, 100 μM AP5, and 100 μM AP5 + 7.78 μM TBOA. **D**. Summary of AP5 effects from 4 animals at different concentrations (30 μM and 100 μM AP5). AP5 alone did not significantly change SWR amplitude, but TBOA still produced significant suppression in the presence of AP5.

Since AP5 did not rescue TBOA-induced SWR suppression, we next tested whether reducing fast AMPA receptor–mediated excitation could preserve SWR activity under TBOA treatment. AMPA antagonist CNQX can eliminate SWRs at a low concentration of 1–2 μM (Fig 5A), about one-tenth of the concentration used for partially blocking AMPA receptors in CA1 neurons [52–53]. To test the effect of CNQX on the TBOA-induced SWR amplitude reduction, we have titrated CNQX in submicromolar concentrations (Fig 5 A–B). SWRs could still be sustained in lower CNQX concentrations (Fig 5A), but the combined application of TBOA and low-dose CNQX exacerbated the TBOA-induced SWR amplitude reduction (Fig 5B).

### H channel blockers did not reverse TBOA effects

Hyperpolarization-activated cyclic nucleotide–gated (HCN) channels are cation channels that open during membrane hyperpolarization and carry the nonselective inward current Ih [57], which contributes to depolarizing the membrane and regulating the resting membrane potential. In principle, reducing Ih could enhance membrane hyperpolarization and thereby counteract the network hyperexcitability induced by TBOA.

To test this possibility, we partially blocked Ih using the HCN channel blocker ZD7288 [58]. Perfusion with 10 µM ZD7288 initially produced a pronounced increase in sharp wave amplitude, consistent with a previous report [59]. However, a complete suppression of sharp waves was observed afterward (Fig 6a). This effect was irreversible upon washout (Fig 6a). Although ZD7288 caused an upward shift in the SWR amplitude distribution, similar to the effects of KCNQ activation and lowering extracellular potassium, it did not rescue the TBOA-induced suppression of SWRs (Fig 6b).

**Fig 6.**
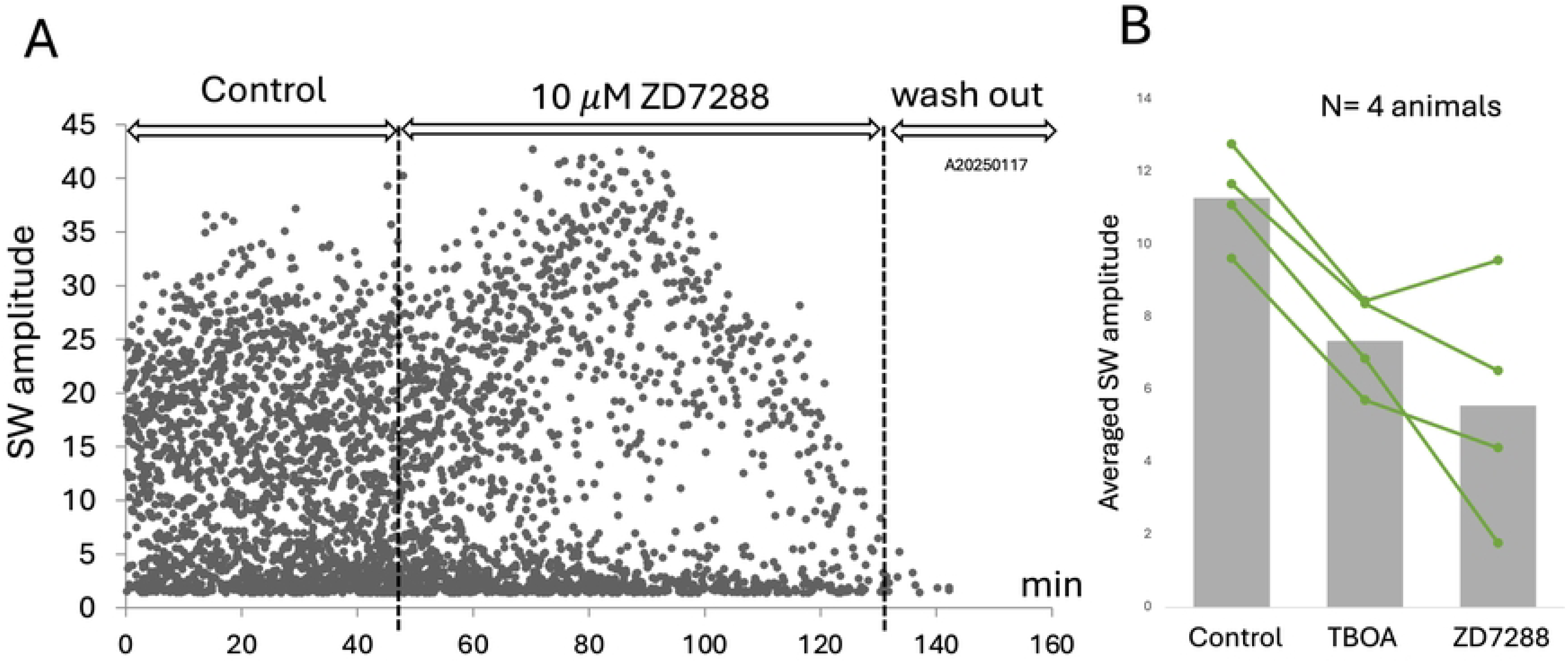
HCN channel blockade by ZD7288 did not rescue TBOA-induced suppression of sharp waves. **A.** Time course of sharp wave (SW) amplitude during perfusion with 10 μM ZD7288 followed by washout. ZD7288 initially increased SWR amplitude, followed by a progressive decline and eventual suppression of SWR activity. The effect was largely irreversible after washout. **B.** Summary of averaged SWR amplitude from 4 animals under control conditions, after TBOA treatment, and during ZD7288 perfusion. Although ZD7288 produced an initial upward shift in SWR amplitude similar to that observed with KCNQ openers and low extracellular potassium, it did not rescue TBOA-induced suppression of SWs and instead led to further reduction of SWR activity.

In conclusion, KCNQ channel activation mitigated TBOA-induced SWR suppression, and this effect was unique to KCNQ activators. Although membrane hyperpolarization can be achieved through multiple mechanisms, including modulation of BK, GIRK, NMDA, and HCN channels, these alternative pathways did not provide similar protection. These results suggest that mild hyperpolarization alone is insufficient; instead, KCNQ-mediated stabilization may be specifically important for preserving organized hippocampal network activity under glutamate dysregulation.

## Discussion

This study yielded two primary findings: (1) TBOA suppressed SWR events in the CA3–CA1 hippocampal network, and (2) KCNQ channel openers partially reversed this suppression and restored SWR activity.

### TBOA suppressed SWR events

TBOA inhibits glutamate transporters, leading to elevated extracellular glutamate [21, 16]. Increased extracellular glutamate is expected to depolarize neurons and increase network excitability. TBOA-induced hyperexcitability has been reported to depend on ongoing neuronal activity [8]. Elevated glutamate can increase spontaneous neuronal firing within the network, and increased firing may further elevate glutamate release, creating a positive feedback loop. This harmful positive feedback has been described as the “vicious cycle of glutamate-mediated hyperexcitability” [8]. Our observation that TBOA suppresses SWRs appears inconsistent with this vicious-cycle model. If network excitability simply increases, one would expect more neurons to participate in each SWR event, producing larger SWR amplitudes rather than smaller ones. One possible explanation for this discrepancy is that elevated extracellular glutamate increases spontaneous neuronal firing but disrupts the temporal coordination required for SWR generation. In this way, glutamate-induced depolarization may shift the network toward disorganized hyperactivity rather than organized population events. SWRs require a large population of neurons to fire within a narrow temporal window; even small shifts in membrane potential that increase background spiking may disrupt this temporal coordination [25, 61]. As a result, network activity becomes more irregular and fragmented, leading to reduced SWR amplitude despite increased overall excitability.

### KCNQ channel activation partially rescues the TBOA effect

Our second primary finding was that TBOA-induced SWR suppression could be partially reversed either by enhancing potassium conductance through KCNQ channel activation (Fig 3) or by lowering extracellular potassium concentration in the perfusate ACSF (Fig 4). KCNQ channels provide a relatively steady subthreshold stabilizing current that may directly counteract TBOA-induced depolarization. The effects of both KCNQ activation and low-potassium ACSF support the interpretation that reducing background firing and improving temporal coordination across the network can help restore the synchronized population activity required for SWR generation.

### Partial rescuing

We noticed that the KCNQ rescuing effect was only partial, as the reduced slope of SWR amplitude declined under TBOA. This suggests that mild hyperpolarization cannot prevent the accumulation of extracellular glutamate. SWRs are large scale population neuronal events, with about ∼5% of CA1 neurons firing at least one spike during each event [25,38, 61]. Each SWR event likely involves substantial synaptic glutamate release, including at Schaffer collateral–CA1 pyramidal neuron synapses and CA1 pyramidal neuron–interneuron synapses [32, 65]. In our *ex vivo* slice system with low cholinergic tone, glutamate production from disorganized spiking is likely small compared with that generated by recurrent SWRs occurring continuously at 0.5–2 Hz. Thus, even if KCNQ openers completely suppress disorganized irregular spiking, glutamate released during SWRs may still accumulate, leading to only partial rescue.

### Negative feedback mitigates the vicious cycle

We speculate a negative-feedback mechanism that may be against the positive-feedback of “vicious cycle,” [8] in the SWR production. During the vicious cycles elevated extracellular glutamate promotes higher network activity which further increases glutamate accumulation (Fig 7 ,left).

**Fig 7.**
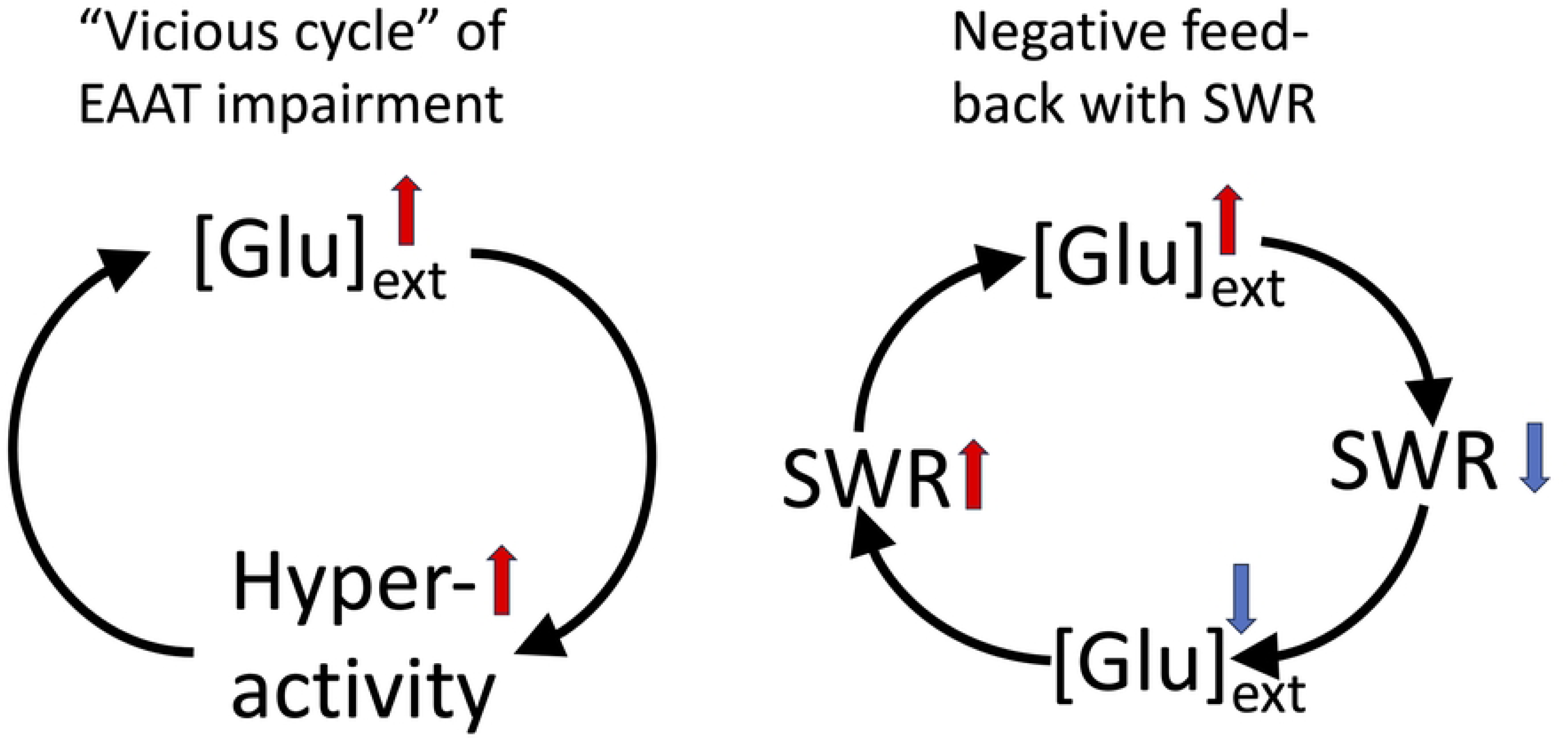
Proposed positive and negative feedback mechanisms during EAAT impairment. *Left panel*: “vicious cycle” induced by EAAT impairment (modified from [8]). Reduced glutamate uptake elevates extracellular glutamate concentration ([Glu]ext), which promotes neuronal hyperactivity and disorganized firing. Hyperactivity further increases glutamate release, producing additional accumulation of extracellular glutamate. Red arrows indicate excitatory or amplifying interactions. *Right panel:* Proposed negative feedback mechanism involving SWRs. SWRs contribute to extracellular glutamate production; however, elevated glutamate suppresses SWR activity. Reduction of SWRs may subsequently decrease SWR-associated glutamate release, partially limiting further glutamate accumulation. Blue arrows indicate reduction of interactions.

In the negative feedback in the SWR production, glutamate-mediated suppression of SWRs would reduce further glutamate release, while the resulting reduction in extracellular glutamate would in turn permit a low level of SWRs to persist (Fig 7 right). Thus, suppression of SWRs may partially counteract the vicious cycle by limiting glutamate accumulation. Consistent with this possibility, we found TBOA never completely abolished SWRs even during many hours of perfusion (data not shown). Further studies are needed to determine whether this proposed negative-feedback mechanism contributes to stabilization of network activity under mild glutamate dysregulation in early AD pathology.

### Uniqueness of KCNQ opener

In our experiments, only voltage gated potassium channel openers can rescue TBOA induced SWR impairment, whereas other potassium channel–based approaches, including BK and GIRK channel openers, failed to restore SWR activity. This result was unexpected because BK and GIRK channel openers are also expected to reduce neuronal excitability, yet neither treatment restored SWR activity under TBOA exposure.

Although this report did not further explore why BK channel and GIRK channel openers were ineffective, their limited effects may reflect mechanisms distinct from KCNQ channel activation. BK channel activation can attenuate inhibitory postsynaptic currents at GABAergic synapses [62], which may disturb the excitation-inhibition balance required for SWR generation. GIRK channels, although hyperpolarizing, are inwardly rectifying K⁺ channels which provide less outward K⁺ current during depolarized network states than KCNQ channels, making them less effective at opposing TBOA-induced depolarization [47]. In fact, GIRK channel activations via 5HT-1A receptors are known to suppress SWRs [64].

HCN channel blockade showed a biphasic effect on SWR activity, initially increasing SWR amplitude before progressively impairing SWR initiation. (Fig 5); NMDA receptor antagonist, does not affect SWRs and show no rescue effects (Fig 5). Generation of SWRs requires delicate balance between excitatory and inhibitory balance [38], pharmacology manipulations to the network may have complex effects on different components for the generation of population activity.

Our results identify a group of compounds capable of reversing amyloid beta–related network impairment and restoring neuronal activity critical for memory consolidation. Indeed, KCNQ channels (M current) underlie are a key factor for the generating of SWRs. SWR is a network activity spontaneously occurring during sleep and quiet restfulness [24, 26, 27], when cholinergic tone is low in the brain. During awake exploratory state, acetylcholine released from basal forebrain attenuates the M-current via muscarinic receptors, leading to the termination of SWRs and transition to the theta oscillation mode.

Cholinergic mediated brain state transition is important for the brain wide activity for consolidation of episodic memory [24,28–31], and memory-guided decision making during awake [63].

Impaired glutamate recycling has been recognized as a major contributor to the synaptic toxicity of soluble amyloid-β, which can disrupt glutamate uptake and elevate extracellular glutamate levels. Our results suggest that such glutamatergic dysregulation may impair coordinated hippocampal network activity by inducing mild depolarization of the neuronal population. Restoring membrane potential balance through potassium-mediated mechanisms may therefore represent a potential strategy to preserve physiological network dynamics under conditions of amyloid-related glutamate dysregulation.

